# Rapid adaptation to a globally introduced virulent pathogen in a keystone species

**DOI:** 10.1101/2024.09.16.613142

**Authors:** Loren Cassin-Sackett, Mirian TN Tsuchiya, Rebecca B. Dikow

## Abstract

Emerging infectious diseases are one of the foremost contemporary threats to biodiversity conservation. Outbreaks of novel pathogens can lead to extinction of host populations, loss of gene flow due to extirpation, and bottlenecks in host populations with surviving individuals. In outbreaks with survivors, pathogens can exert strong selection on hosts, in some cases leading to the evolution of resistance or tolerance in the host population. The pathogen causing sylvatic plague, *Yersinia pestis*, was introduced to North America in the early 20^th^ century and caused rapid population declines in prairie dogs (genus *Cynomys*), which experience >95% mortality during epizootics. Recently, survival from plague has been documented in a small number of black-tailed prairie dogs (*C. ludovicianus*) in natural populations in Colorado (USA). We performed whole-genome sequencing on 7 individuals from 3 colonies that survived infection with plague and 7 individuals from the same colonies that likely died during a plague epizootic. Using genome-wide association tests, F_ST_ outlier tests, and other inferences of selection, we detected SNPs on 5 scaffolds that were strongly associated with survivorship from plague in the wild. Some genes associated with these scaffolds also differ in humans that survived versus died in the plague pandemic in London, UK, suggesting conservation of gene function across taxonomically diverse lineages. Understanding the genetic basis of immunity can enable genetically-informed management actions such as targeted relocation to protect prairie dogs and the species that rely on them. More generally, understanding how rapid adaptation to pathogens occurs can help us predict the time frame and spatial scale at which adaptation may occur, during which other interventions are needed.

**Significance Statement:** Emerging infectious diseases are one of the foremost threats to global biodiversity, causing extinctions and population crashes on all continents. Introduced pathogens can exert strong selection on hosts for the evolution of tolerance or resistance, yet these evolutionary events are rare and it remains challenging to identify and sample both immune and susceptible individuals during an epizootic. This study leverages one of the only documented examples of prairie dogs surviving infection from introduced sylvatic plague in nature and compares their genomes to those of individuals that perished. We find strong signatures of selection in a small number of immune and non-immune genes, one of which has been implicated in survival from plague in humans. These findings suggest that adaptation to novel pathogens may occur via a combination of conserved genes and the co-opting of genes outside of classical immune pathways. Finally, it provides evidence that in native species with sufficient standing genetic variation, there is potential for adaptation to introduced pathogens.

## Introduction

Species across the globe are increasingly confronted with strong novel selection pressures that threaten them with extinction if populations do not adapt (Thomas et al. 2004; Pearson et al. 2014; McLaughlin et al. 2002; Uthicke, Momigliano, and Fabricius 2013; Pillet et al. 2022; Gómez et al. 2015; Otto 2018). Anthropogenic change has increased the rate of climate warming and extreme events, habitat alteration, introduced species, and emerging infectious diseases (Trenberth, Fasullo, and Shepherd 2015; Cook et al. 2016; Fahrig 2007; Vitousek et al. 1997; Benning et al. 2002), which can act in concert to exert multifarious novel selection. In particular, climate change facilitates the range expansion of some introduced species (Atkinson et al. 2014; Walther et al. 2009; Blumenthal et al. 2016), exerting selection on native species in the invaded range (Dufour, Herrel, and Losos 2017; Fazlioglu and Chen 2020; Strauss, Lau, and Carroll 2006) that can lead to adaptation (Woodworth et al. 2005) or extinction (Savidge 1987; Freed, Cann, and Bodner 2008; Dangremond, Pardini, and Knight 2010).

Introduced pathogens are a primary driver of global population declines of native species (e.g., canine distemper in carnivores (Panzera et al. 2015; Seimon et al. 2013; Deem et al. 2000; Roelke-Parker et al. 1996), avian malaria in Hawaiian honeycreepers (Warner 1968; Cassin-Sackett et al. 2021), white-nose syndrome in bats (Drees et al. 2017; Frick et al. 2010; Zukal et al. 2016), and chytridiomycosis in amphibians (Lips et al. 2006; Scheele et al. 2019; O’Hanlon et al. 2018)), and they also threaten crops (Ali et al. 2014; Callaway 2016) and livestock (Zimpel et al. 2020; Kitching, Thrusfield, and Taylor 2006). Although some species have demonstrated rapid adaptation (Bonneaud et al. 2011; Best and Kerr 2000; Woodworth et al. 2005; Epstein et al. 2016; Auteri and Knowles 2020), we lag in our understanding of which types of genes underlie adaptation to novel pathogens in nature (Schiebelhut, Puritz, and Dawson 2018; Ebert and Fields 2020). As globalization intensifies, there is a pressing need to understand the genetic basis of recently evolved immunity to introduced pathogens and to translate this information to practical solutions such as assisted gene flow, targeted relocation, or reintroductions that maximize genetic variation in natural populations.

Understanding the genetic basis of rapid adaptation to a selection pressure introduced on a known time scale can illuminate under what circumstances naive hosts may possess the genetic variation that facilitates adaptation. When adaptation to pathogens occurs on very short time scales, the genomic changes may be costly (e.g., sickled red blood cells in humans infected with *Plasmodium falciparum* (Taylor, Cerami, and Fairhurst 2013; Lansche et al. 2018)), variable across populations (Gignoux-Wolfsohn et al. 2021; Elfekih et al. 2022; Kuttiyarthu Veetil et al. 2024), and may invoke non-immune as well as immune genes (Cassin-Sackett, Callicrate, and Fleischer 2019; Elfekih et al. 2022), each with a different evolutionary prognosis for the host. Therefore, identifying which types of genes are involved in the rapid evolution of host immunity to novel pathogens can illuminate the most effective strategies to prevent widespread extinctions.

The bacterial pathogen that caused the Black Death in humans, *Yersinia pestis*, has been introduced around the world and continues to cause deadly outbreaks in humans (Andrianaivoarimanana et al. 2019) and wildlife (Tang et al. 2022; Bevins et al. 2021). The pathogen first arrived in North America via rat fleas stowed aboard ships that docked in California in 1900 (Eskey and Haas 1939), with likely additional introductions to Seattle in 1907 and Los Angeles in 1908 (Adjemian et al. 2007), and it subsequently spread eastward to the 100^th^ meridian (Adjemian et al. 2007). *Yersinia pestis* causes sylvatic plague in numerous species of mammals, most notably ground squirrels (Wherry 1908; Williams, Moussa, and Cavanaugh 1979) and the critically endangered black-footed ferret (Matchett et al. 2010). Species such as the highly social prairie dogs (genus *Cynomys*) are extremely susceptible, and their populations experience stunning declines during epizootics: Plague causes 95-100% mortality in prairie dog populations (Ecke and Johnson 1952; Cully et al. 1997; Lechleitner et al. 1968; Pauli et al. 2006), and die-offs can happen within a few weeks (Lechleitner et al. 1968).

Although plague causes local extirpations, populations of prairie dogs are often recolonized from multiple source populations, leading to a retention of genetic variation even in fragmented landscapes (Sackett, Collinge, and Martin 2013a). The strong selection exerted by plague and the potential for maintenance of genetic variation suggests prairie dogs may possess standing variation for immunity to plague; indeed, survival from plague in natural populations of prairie dogs has been documented (Cully et al. 1997; Pauli et al. 2006; Sackett, Collinge, and Martin 2013a), although survival in the wild appears rare. Experimental plague infections show higher survivorship in prairie dogs from geographic regions where plague was introduced earlier, relative to prairie dogs from regions where plague arrived later (Rocke et al. 2012). Collectively, this evidence suggests there may be a heritable genetic basis to survivorship from plague.

Here, we capitalize on a plague epizootic in natural populations of black-tailed prairie dogs (*C. ludovicianus*) during which we sampled 7 individuals that produced plague antibodies, 6 of which we recaptured the following year, indicating survivorship from plague. We use whole-genome sequencing to conduct a set of genome-wide association tests in conjunction with tests for sites under selection to search for evidence of evolved immunity in these prairie dogs that survived infection with *Y. pestis*.

## Methods

### Field Sampling and Plague Epizootic

One individual sampled in 2017 was selected for whole-genome assembly based on its high quantity and quality of DNA and its low individual heterozygosity (because low-heterozygosity individuals have traditionally been more straightforward to assemble). Ear tissue was collected and used for 24 parallel extractions using the Qiagen DNeasy Blood and Tissue kit; multiple extractions from the same individual ensured sufficient DNA quantity. Low heterozygosity at microsatellite loci (0.438) was confirmed by genotyping the individual at 16 nuclear loci (as in (Sackett, Collinge, and Martin 2013a)).

From 2006 - 2009, a plague epizootic spread through Boulder County, Colorado (USA) in the sites of a long-term prairie dog ecology study (W. Johnson and Collinge 2004; Collinge et al. 2005; Snäll et al. 2008; Brinkerhoff et al. 2011; Jones, Knight, and Martin 2010; Bai et al. 2008; Brinkerhoff et al. 2010; Sackett et al. 2012; Sackett, Collinge, and Martin 2013a). The epizootic extirpated nearly all colonies, but a small number of individuals (<20) in each of three colonies remained (Fig 1a). We live-trapped prairie dogs in these surviving colonies during and after the epizootic (scientific collection license #08TR2012). Capture methods have been described in detail elsewhere (Sackett, Collinge, and Martin 2013a); briefly, we placed 1 - 4 Tomahawk traps near the entrances of active burrows, pre-baited traps for at least 5 days, and set traps for one to two weeks in an attempt to maximize the number of individuals captured.

**Fig 1.**
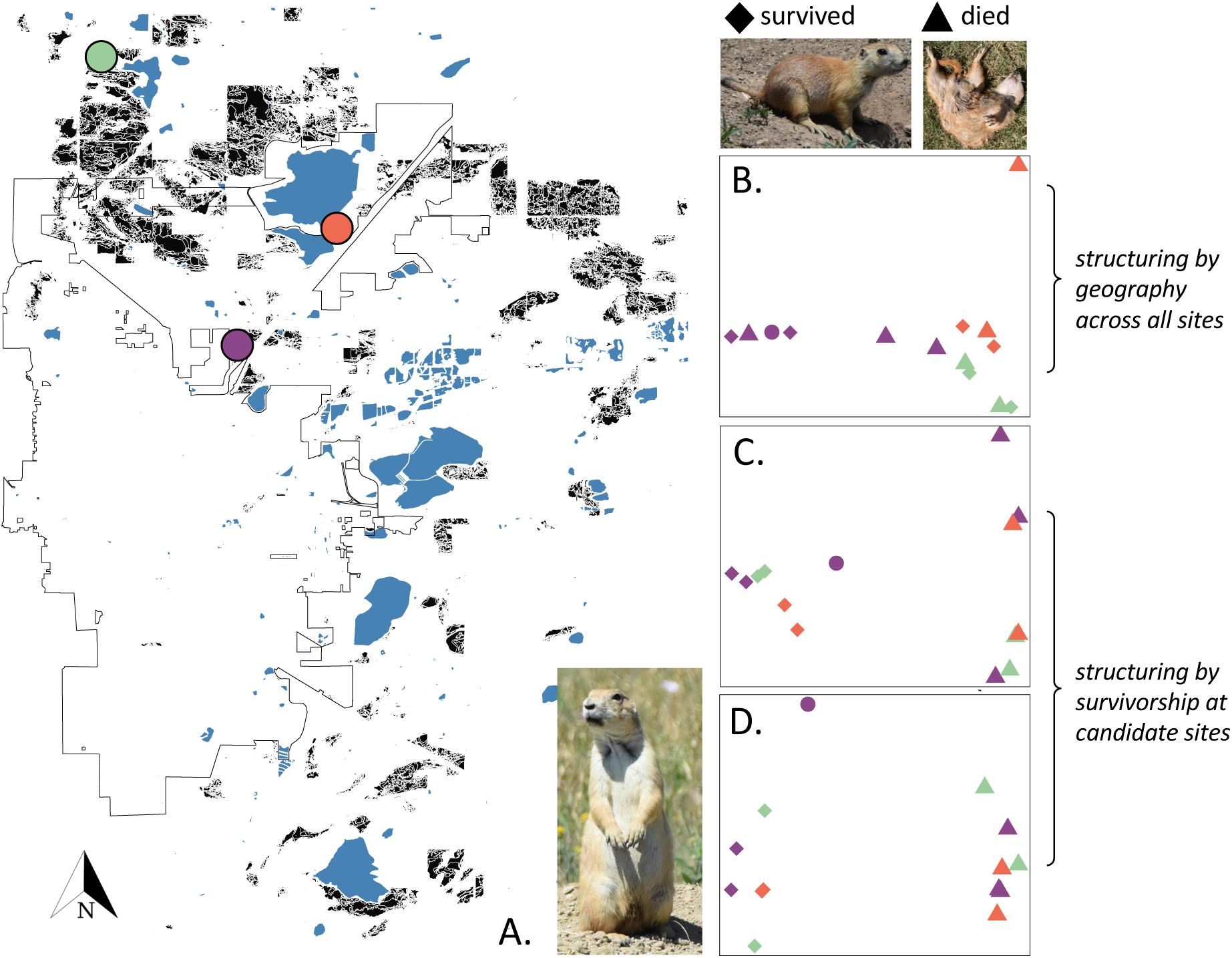
A. Map of prairie dog colonies (black shapes) surrounding the city of Boulder, CO (dark gray outline), with our sampling sites shown in colored circles (green = Beech, coral = Reservoir, purple = Belgrove). Inset: black-tailed prairie dog from near Reservoir. B. Principal Component Analysis (PCA) of ∼7.9 million SNPs genotyped in 14 individuals from three colonies. Genotypes separate somewhat by geography (colors as in part A), but not by survivorship phenotype (shapes: diamonds=survivors, circle=antibodies/not recaptured, triangles=fatalities). C. PCA of 1334 candidate SNPs at moderate threshold; genotypes do not segregate by sampling location (colors) but do segregate by phenotype (shapes). D. PCA of 154 candidate sites at stringent threshold; clustering is by resistance phenotype. Antibody individual clusters with survivors at candidate loci (C and D).

Captured animals were anesthetized with isoflurane for the collection of tissue and blood. A one cm piece of tissue was trimmed from the outer ear, and 0.5mL blood was collected from the femoral vein. Tissue was stored in a solution of 20% dimethyl sulfoxide in EDTA to preserve the DNA (Seutin, White, and Boag 1991). Blood was spread evenly on a Nobuto strip (Advantec MFS, Dublin, CA), dried, and sent to the CDC for a hemagglutinin assay to detect antibodies to *Yersinia pestis* (May Chu, n.d.). Demographic information was recorded and the animal was subsequently returned to its trap, placed in the shade and allowed to recover from anesthesia prior to release at its capture location. All protocols were approved by the University of Colorado’s Institutional Animal Care and Use Committee (Protocol # 08-07-SAC-01).

### De novo Reference Genome Assembly

Although there currently exists a reference genome for another species in the genus (Tsuchiya, Dikow, and Cassin-Sackett 2020), there is considerable variation in karyotype within the genus (Nadler, Hoffmann, and Pizzimenti 1971; Pizzimenti 1976) that warranted a new assembly for black-tailed prairie dogs. We assembled the genome using the same procedures as (Tsuchiya, Dikow, and Cassin-Sackett 2020). Briefly, libraries were prepared from replicate DNA extractions and samples were sequenced to 20x on a PacBio Sequel and to 80x on an Illumina HiSeq 4000 (150-bp paired-end reads) at Duke University’s Sequencing and Genomic Technologies Shared Resource core facility. The genome was assembled *de novo* by combining long and short reads in MaSurCA v. 3.8.1 (Zimin et al. 2017), and further scaffolding was performed using SSPACE-LongRead (Boetzer and Pirovano 2014). Gaps were filled using PBJelly (English et al. 2012), and the genome was polished in Pilon (Walker et al. 2014). We used Kraken (Wood, Lu, and Langmead 2019) to remove scaffolds classified as bacteria. We used Benchmarking Universal Single-Copy Orthologs (BUSCO v. 3.0.2, (Simão et al. 2015) to assess the completeness of the assembly by comparing it to the mammalia_odb9 gene database (Zdobnov et al. 2017).

### Whole-Genome Resequencing of Plague-Infected Animals

We selected 14 individuals for whole-genome sequencing based on survivorship outcome (N=7 survivors, N=7 presumed fatalities). Samples were matched by sex, age, and source colony. DNA was extracted using a Qiagen DNeasy Blood and Tissue Kit following the manufacturer’s protocol and genomic DNA was sent to Novogene (Sacramento, CA) for library preparation and whole-genome sequencing. Genomes were sequenced to a depth of 30X on an Illumina HiSeq X using 150bp paired-end reads. Demultiplexed sequences were quality filtered for a minimum SNP quality score of 20 and aligned to the black-tailed prairie dog reference genome using Bowtie2 (Langmead and Salzberg 2012), and we called variants on each individual separately using the GATK v4 HaplotypeCaller (McKenna et al. 2010; Van der Auwera and O’Connor 2020) Next, we excluded all SNPs with coverage depth < 8 and > mean*2SD; the latter was designed to remove SNPs that were found in paralogs, which can be misinterpreted due to inflated diversity at those sites. Finally, we excluded SNPs that were not genotyped in all individuals, resulting in a final dataset containing 7,863,840 SNPs and 61,388 MNPs. All further analyses using this dataset, described below, are detailed at https://github.com/CassinSackett/PlagueResistance.

### Association Test

Prior to the association test, we verified that sampled individuals were not siblings or parents/offspring by estimating relatedness among individuals in vcftools (Danecek et al. 2011), and we estimated genetic structure both by calculating Weir and Cockerham’s F_ST_ in vcftools and by conducting an analysis of molecular variance (AMOVA) in the poppr v2.9.4 package (Kamvar, Brooks, and Grünwald 2015) for R v 4.3.0 (R Core Team 2023) using the ade4 implementation (Dray and Dufour 2007; Excoffier, Smouse, and Quattro 1992). We visualized genetic structure by plotting the first two axes of a Principal Component Analysis (PCA).

Due to the statistical uncertainty inherent in small sample sizes and the number of false positives that can arise from testing >7 million SNPs (Hoban et al. 2016), we aimed not to detect all significant SNPs but rather to focus on a smaller number of stronger candidates. To do so, we performed multiple genome-wide association tests to test for alleles associated with survivorship from plague and examined loci that were significant across multiple tests. We performed tests in plink v1.9 (Purcell et al. 2007; Chang et al. 2015) and plink2 (www.cog-genomics.org/plink/2.0/), using a combination of chi-square (plink v1.9, --assoc), Cochran-Armitage trend tests (plink v1.9, --model trend (Steiß et al. 2012)), and general linear models (plink2, --glm) and evaluating significance based on genomic control-corrected p-values (Devlin and Roeder 1999) and Landcaster’s mid-P correction to the Fisher’s exact test (Biddle and Morris 2011). With the --assoc test in plink v1.9 we used the --ci option (Steiß et al. 2012) to generate odds ratios with 95% confidence intervals for each SNP, and we examined both high and low odds ratios because our sample size was balanced between survivors and fatalities, leading to some sites with a minor allele found primarily in survivors and other sites with a minor allele found primarily in fatalities. We also examined loci with significant p-values for which odds ratios could not be calculated due to an alternative allele frequency of 0 in the survivor group (i.e., when all survivors had the same allele as the reference). We performed most of the plink v1.9 tests two different ways - (1) including the individual that produced antibodies but was not recaptured as a survivor and (2) excluding this individual from the test.

In plink2, we compared a model with no covariates to two models accounting for genetic structure in the sample (with the first 3 axes of a PCA, and with the first 4 axes of a PCA). We did not include additional axes as covariates due to the small number of samples. We examined the lambda inflation factor for each test to select the best parameter set that avoided overfitting the model. We selected the model with no covariates because lambda was closest to 1 (0.963). This decision was justified by the close sampling location of sampled individuals (Fig 1a) and supported by an AMOVA (Excoffier, Smouse, and Quattro 1992) (using the poppr.amova function in the poppr package for R; (Kamvar, Tabima, and Grünwald 2014), which found less than 3% of genomic variation to be partitioned among sampling locations. However, because all three plink2 models contained the same set of 193 significant SNPs, and the PC1-4 model was the only one able to distinguish among this set of SNPs, we selected the 20 most-significant SNPs from this model to include in our “stringent threshold” set of candidates (see results). Finally, for each test, we collated the most-significant SNPs in each test; for example, genomic-control adjusted p<0.001 or fisher mid-p < 1e-4). We chose to evaluate the most-significant SNPs in each model, rather than setting a consistent threshold across all models, due to differences in power among models. We selected our final set of 65 candidate SNPs as those that were common across multiple tests or found in clusters of >=10 SNPs on a scaffold. We also generated a list of SNPs that were significant at a slightly less stringent threshold (Table Sx). A summary of models and significance thresholds is provided in Table 1.

**Table 1.** List of models used to infer sites under selection and significance threshold used to identify candidates. Models were tested that compared the 7 presumed fatalities with only the known survivors (N=6 recaptured the year after their first detection with antibodies), with the non-recaptured individual that produced antibodies excluded (e.g., “6 controls 7 cases”), and another set of models was conducted with all antibody-producing individuals included as survivors (e.g., “7 controls 7 cases”).

| Test | Parameters | Samples | moderate threshold for evaluation | # SNPs | stringent threshold for evaluation | # SNPs | most stringent threshold for evaluation | # SNPs |
| --- | --- | --- | --- | --- | --- | --- | --- | --- |
| plink1.9 | --assoc | 6 controls<br>7 cases | Odds Ratio > 40 | 140 | Odds Ratio > 65 | 28 | Odds Ratio > 65 | 28 |
| plink1.9 | --assoc | 7 controls<br>7 cases | Odds Ratio<br>> 45 | 73 | Odds Ratio<br>> 77 | 16 | Odds Ratio<br>> 169 | 2 |
| plink1.9 | --assoc | 6 controls<br>7 cases | Odds Ratio<br>< 0.1 and<br>frequency<br>diff > 0.8 | 51 | Odds Ratio<br>= 0 and<br>FMIN_U ><br>0.8 | 20 | Odds Ratio<br>= 0 and<br>freq-diff ><br>0.9 | 3 |
| plink1.9 | --assoc | 7 controls<br>7 cases | Odds Ratio<br>< 0.1 and<br>frequency<br>diff > 0.8 | 26 | Odds Ratio<br>= 0 and<br>FMIN_U ><br>0.8 | 10 | Odds Ratio<br>= 0 and<br>freq-diff ><br>0.9 | 1 |
| plink1.9 | --assoc --<br>adjust | 6 controls<br>7 cases | genomic-control p <<br>5.0x10e-4 | 359 | genomic-control p <<br>1.05 x10e-4 | 20 | genomic-control p <<br>2.6 x10e-5 | 6 |
| plink1.9 | --assoc --<br>adjust | 7 controls<br>7 cases | genomic-control p <<br>7.4x10e-4 | 214 | genomic-control p <<br>8.75 x10e-05 | 11 | genomic-control p <<br>2.5 x10e-05 | 1 |
| plink1.9 | --model<br>trend --<br>adjust | 6 controls<br>7 cases | genomic-control p <<br>3.0x10e-3 | 193 | N/A |  | N/A |  |
| plink1.9 | --model<br>trend --<br>adjust | 7 controls<br>7 cases | genomic-control p <<br>5.18 x10e-3 | 73 | genomic-control p <<br>2.6 x10e-3 | 42<sup>s</sup> | N/A |  |
| plink1.9 | --assoc --<br>fisher-midp | 6 controls<br>7 cases | fisher mid-p<br>adjusted p <<br>3.6 x10e-5 | 186 | fisher mid-p<br>adjusted p<br>< 6.3 x10e-6 | 20 | fisher mid-p adjusted<br>p < 1.4<br>x10e-6 | 6 |
| plink1.9 | --assoc --<br>fisher-midp | 7 controls<br>7 cases | fisher mid-p<br>adjusted p <<br>7.7 x10e-05 | 214 | fisher mid-p<br>adjusted p<br>< 5.0 x10e-06 | 31 | fisher mid-p adjusted<br>p < 3.8<br>x10e-07 | 1 |
| plink1.9 | --model<br>trend --<br>fisher-midp<br>(recessive<br>model) | 7 controls<br>7 cases | fisher mid-p<br>adjusted p <<br>2.4 x10e-03 | 37 | N/A |  | N/A |  |
| plink1.9 | --model<br>trend --<br>fisher-midp<br>(dominance<br>model) | 7 controls<br>7 cases | fisher mid-p<br>adjusted p <<br>3.00x10-04 | 419 | N/A |  | N/A |  |
| plink1.9 | --model<br>trend --<br>fisher-midp<br>(genotype<br>model) | 7 controls<br>7 cases | fisher mid-p<br>adjusted p <<br>3.00-04 | 679 | N/A |  | N/A |  |
| plink2 | no<br>covariates | 7 controls<br>7 cases | genomic<br>control p <<br>0.01088 | 193 | N/A |  | N/A |  |
| plink2 | PC 1 - 3 covariates | 7 controls<br>7 cases | genomic control $p < 0.026$ | 213 | N/A | | N/A | |
| plink2 | PC 1 - 4 covariates | 7 controls<br>7 cases | genomic control $p < 0.034$ | 213 | genomic control $p < 0.0335$ | 20<sup>#</sup> | N/A | |
| BayPass | Core model | 6 survivors<br>7 fatalities | $-\log(p) > 6$ | 112 | $-\log(p) > 6.6$ | 41 | $-\log(p) > 7.5$ | 3 |
| $F_{ST}$ | Weir & Cockerham $MAC=2$ | 6 survivors<br>7 fatalities | $F_{ST} > 0.75$ | 160 | $F_{ST} > 0.8$ | 54 | $F_{ST} > 0.9$ | 6 |
| $F_{ST}10kb$ | Weighted $F_{ST}$ | 7 survivors<br>7 fatalities | $F_{ST} > 0.5 + \geq 10$ SNPs in window | 149 windows | $F_{ST} > 0.6 + \geq 10$ SNPs in window | 26 windows | $F_{ST} > 0.64 + \geq 10$ SNPs in window | 4 |

N/A in the “stringent threshold” column means there were no SNPs at a lower significance threshold than the moderate threshold
<sup>§</sup> trend model did not recover any SNPs that were also significant in other GWAS models
<sup>#</sup> none of the plink2 SNPs were recovered from other GWAS

### Tests of selection

First, we used BayPass v2.4 (Gautier 2015) to test for sites under selection under the model articulated by Coop et al. (2010) and Gunther & Coop (2013). We generated a list of genotyped sites in the vcf file, converted our vcf file to BayeScan format using the publicly available vcf2bayescan.pl perl script (https://github.com/santiagosnchez/vcf2bayescan) and then converted the resulting BayeScan file to BayPass format using the python script geste2baypass.py in geste2baypass (https://github.com/CoBiG2/RAD_Tools/). We used the default settings and ran the standard model with no covariates, as our two groups were from the same sites. We examined the distribution of -log(10) p values and selected the sites in the far right-most tail of the distribution (e.g., corresponding to 0.0005% of sites (-log(10) p > 6.6) for the stringent threshold; Supplementary Figure 1).

Next, we searched for outlier loci between survivors (the 6 antibody-producing individuals that survived) and fatalities (the 7 individuals in very poor body condition that were not recaptured; see Results) in two ways. We examined the distribution of per-site F_ST_ values using the Weir and Cockerham (1984) estimator implemented in vcftools, and selected for follow-up the 54 SNPs that were in the far right tail of the distribution (< 0.0007% of sites; Supplementary Figure 2; F_ST_>0.8). We then calculated F_ST_ in 50kb windows in vcftools, selecting the top 0.05% most differentiated windows (51659 total windows * 0.0005 = 26 high-F_ST_ windows), and removing windows that had <30 SNPs in the window.

To visualize clustering of individuals, we coded genotypes as 0, 1 or 2 in vcftools, where the number corresponds to the number of non-reference alleles, and used the genotypes at candidate loci to generate a heatmap of genotypes of fatalities and survivors. To do so, we used the *dist* function in the stats package for R to calculate manhattan distances and then the *hclust* function to cluster genotypes based on similarity. We visualized this heatmap with the gplots package (Warns et al. 2022) using a modified RColorBrewer (Neuwirth 2022) palette.

Finally, we aimed to identify sites that not only differed between survivors and fatalities, but alleles that were either unique or more common in survivors relative to the reference genome, which we assume to represent a susceptible individual. Thus, we selected for follow-up candidate sites that contained at least one alternate allele in each survivor but only reference alleles in all fatalities. We calculated linkage disequilibrium on a per-scaffold basis for scaffolds that had multiple candidate sites at the moderate or stringent evaluation threshold.

### Linkage Disequilibrium

For each scaffold, we calculated linkage disequilibrium (LD) in plink v1.9 setting a minimum output threshold of r2=0.1 and up to 50 SNPs per window; all other settings were the defaults. Next, we calculated the mean r2 in 1000bp bins and examined the rate of LD decay. The size of the LD block at which LD dropped below half its maximum value in 1000bp bins was determined for each scaffold using a custom R script. We compared the size of LD blocks between candidate and non-candidate scaffolds using a randomization test implemented with a custom R script.

### Candidate Gene Identification

Our general approach for identifying candidates, aiming both to reduce false positives (Cornetti and Tschirren 2020) and identify likely causal variants, was: (1) Start with SNPs recovered in multiple tests, (2) omit SNPs that were the only significant SNP on the scaffold, (3) examine SNPs that contained higher frequencies of alternate alleles in the survivors. For each candidate SNP, we used bedtools to generate a bed file containing regions 500bp upstream and downstream of the SNP and to extract the sequence of that 1000bp region from our reference genome. We then performed a blastn search against the NCBI nucleotide database (updated locally 12/15/23) and examined matches with E values < 1e-40. For all SNPs with no blast matches, we extracted longer sequences in a stepwise manner (2000bp regions, then 3000bp regions, etc.). We inferred the likely function of candidate genes using genecards.org.

## Results

Capture rates in recolonized colonies the year after plague extirpation (or near extirpation) ranged from 7 - 15 (Sackett, Collinge, and Martin 2013a). Seven prairie dogs from three colonies tested positive for antibodies to the *Y. pestis* F1 antigen (Sackett, Collinge, and Martin 2013a), a finding that has been previously reported only a small number of times (Cully et al. 1997; Pauli et al. 2006). Antibodies have not been detected in live captures of other prairie dogs in Boulder County despite testing at least 2,336 samples over 7 years (Colman et al. 2021; Sackett, Collinge, and Martin 2013a). Six of those antibody-producing animals were recaptured the following year; all still had detectable, but lower, antibody titers the following year. The production of antibodies and survival to recapture (N=6) was used as evidence that prairie dogs were resistant or tolerant to plague (i.e., they were exposed to plague and survived). The seventh individual is strongly suspected to also be resistant to plague, given the extreme rarity of the production of antibodies in prairie dogs. Among the individuals not producing plague antibodies, we chose to sequence the individuals with the worst body condition, including three with symptoms suggestive of plague infection (e.g., swollen lymph nodes or other body parts, bloody eyes, numerous patches of bare skin with missing fur, abnormal body odors). These symptoms were not detected during the absence of plague in any of the 24 colonies that were part of the long-term study sites.

### Genome Assembly

The assembly was 2.56 Gb, similar to the congener Gunnison’s prairie dog (Tsuchiya, Dikow, and Cassin-Sackett 2020), and contained 7147 scaffolds, with a scaffold L50 of 708 and scaffold N50 of 1,032,944 bp. Final coverage was 54x with Illumina and13x with PacBio. We recovered 3,862 (94%) complete and 127 (3.1%) fragmented BUSCOs out of 4,104 mammalian orthologs searched.

### Genetic Structure

Four pairs of individuals, all within one colony (17A), had relatedness coefficients between 0.057 - 0.086, consistent with first cousin once-removed or similar relationships. Because there was no consistent pattern of survival among these individuals (e.g., survivors were related to both other survivors and to fatalities), they were all retained in the analyses. Across all ∼7.9 million SNPs, there was some structuring by sampling location (Figure 1b), but <3% of genetic variance was partitioned among populations; most variation was among individuals. Mean Weir & Cockerham’s F_ST_ across the three colonies was 0.020 and weighted F_ST_ was 0.028. When examining 1334 SNPs defined as candidates at a moderate significance threshold (Figure 1c) and 154 SNPs defined as candidates at a stringent threshold (Figure 1d; see “*Candidate loci*” below), structuring by population disappeared and individuals instead clustered by survivorship phenotype (survivors at left denoted by circle and diamonds; fatalities at right denoted by triangles). Structuring at a smaller number of candidate sites (namely, the 20 “top” and 21 “secondary” candidates; see below) is not displayed due to complete overlap in PC scores among some individuals. Mean Weir & Cockerham’s F_ST_ across the three colonies at the 41 top and secondary candidates was -0.142 and weighted FST was -0.149; Weir & Cockerham’s estimates of F_ST_ among colonies were negative at all 41 candidate sites.

### Association Test

At the stringent threshold for evaluation (Table 1), we recovered 154 significant SNPs across all models (Supplementary Table S2), with 64 SNPs being shared across multiple models (Supplementary Table S3). Significant SNPs were primarily located in clusters on 4 scaffolds (scaffold 22, scaffold 375, scaffold 443, and scaffold 519). Significant SNPs that were not common across >2 models were often recovered by multiple models at the less stringent (“moderate”) significance threshold (Supplementary Table S1). At the moderate significance threshold, there were also large numbers of SNPs on 4 other scaffolds (scaffold582, scaffold950, scaffold1325, scaffold1968).

Most tests performed in plink v1.9 each yielded 10-42 SNPs associated with survivorship phenotype at the “stringent significance threshold” for each model (Table 1; Supplementary Table S2), but some models (e.g., the dominance model and genotype model) did not have enough statistical resolution to distinguish differences in significance among the large number of SNPs that were recovered at the “moderate threshold” (Table 1), which recovered 419 and 679 significant SNPs, respectively (p=2.9×10^-4^ in both models). In the models with the moderate threshold (Supplementary Table S1), 1344 SNPs were significantly associated with survivorship phenotype. These SNPs were included in the set of “candidate SNPs” used to conduct diversity analyses, but were not included in the blast or LD analyses and were not considered among the top candidates for follow-up. A PCA at these 1334 moderate candidate SNPs separated individuals by phenotype but not by sampling location (Fig 1c).

Each of the three plink2 models that differed in the numbers of covariates yielded the same 193 SNPs at the moderate threshold, with the PC1-3 and PC1-4 models recovering an additional 20 SNPs. Although the p-values were higher in the PC1-4 model, this was the only model that could differentiate 20 of the top SNPs at a lower significance threshold (Table 1).

The --assoc test in plink1.9 recovered 30 SNPs displaying extremely high odds ratios (OR>65); 27 of these SNPs were found in clusters on 4 scaffolds (scaffold214, scaffold375, scaffold519, scaffold654). In addition to the 30 SNPs with the highest odds ratios, we identified 17 SNPs with allele frequency differences between survivors and fatalities of at least 0.9 and an odds ratio of 0 (and 1 with an odds ratio that could not be computed due to the major allele being fixed in survivors). These 18 SNPs were found on two of the same scaffolds (scaffold214, scaffold519) and on one additional scaffold (scaffold443).

### Tests of selection

BayPass identified a total of 42 sites across 14 scaffolds as being under selection (log10(p) > 6.6). Of these sites, the majority were found in clusters on 7 scaffolds (7 significant SNPs were the only significant SNP on a scaffold). Six of these scaffolds corresponded to scaffolds containing SNPs significantly associated with survivorship or SNPs that were highly differentiated (by F_ST_) between survivors and fatalities.

The distribution of Weir & Cockerham’s F_ST_ values was highly skewed, with a long right tail of extremely high F_ST_ values. We detected 54 SNPs on 6 scaffolds with F_ST_ > 0.820. In our windowed analysis, we detected 21 windows of high F_ST_ across 19 scaffolds, only 3 of which were represented in the set of candidate high-F_ST_ single SNPs (but 3 other scaffolds were detected in the BayPass analysis).

### Linkage Disequilibrium

Linkage disequilibrium remained high across the scaffold for candidate scaffolds (Fig 2f, 3f, Supplementary figures S3 – S5), and LD blocks on candidate scaffolds were larger than on non-candidate scaffolds (p=0.004, Supplementary Fig. S6). This difference was primarily driven by two scaffolds: scaffold 22 contained LD blocks with r2=0.5 across 309,000 bp (Supplementary Fig. S1, S6), and scaffold 519 contained LD blocks with r2=0.5 across 407,000 bp (Fig. 3f, Supplementary Fig. S6).

**Figure 2.**
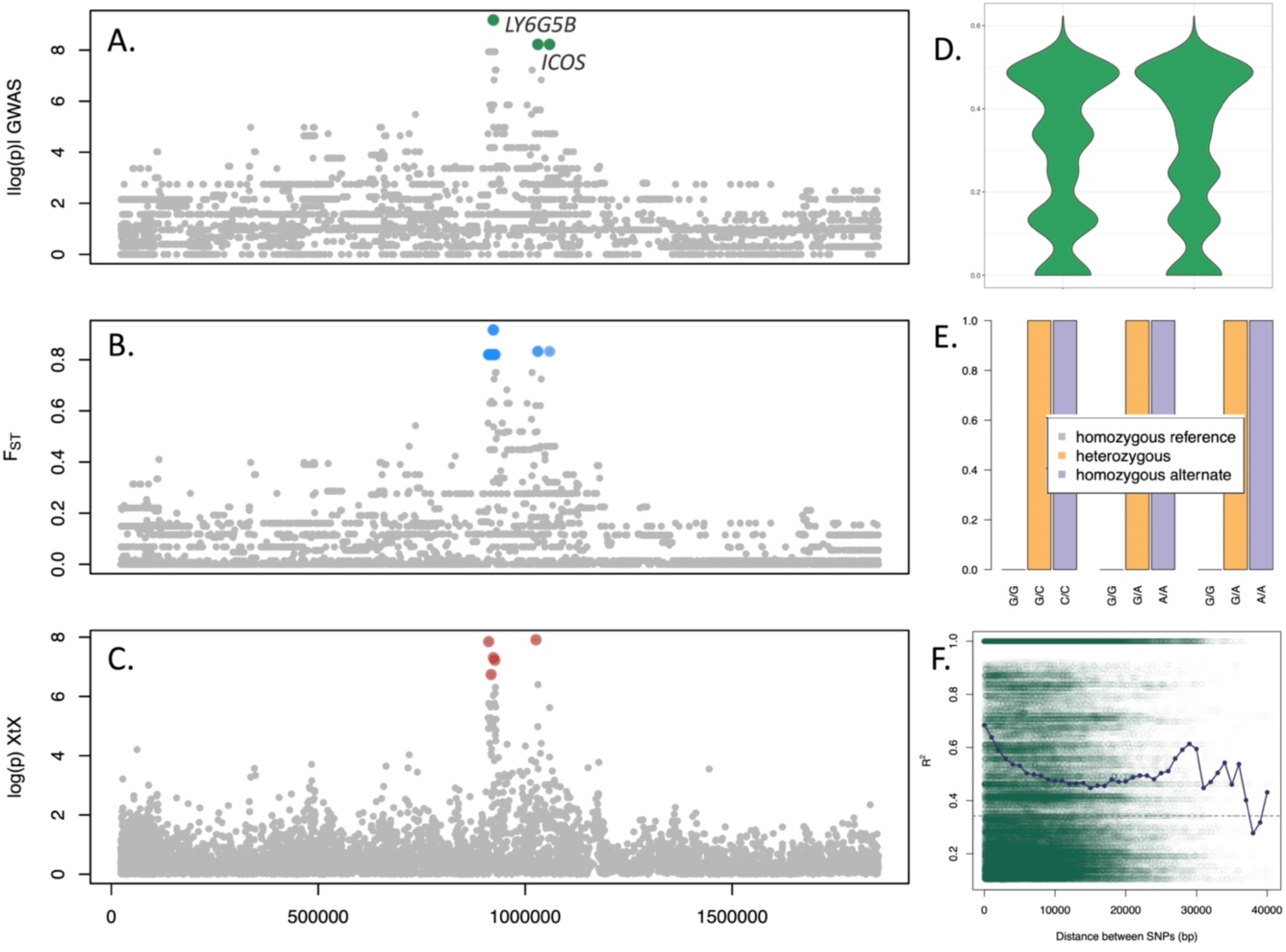
Candidate loci on Scaffold 443. A: -log(p) using genomic control adjusted p-values on a test in plink v1.9 for association between survivorship and genotype. B: F_ST_ between survivors and fatalities. C: - log(p) of the XtX selection statistic calculated in BayPass. D: Heterozygosity on this scaffold in fatalities (left) and survivors (right). E: Percent survivorship for each genotype at the three candidate sites on scaffold 443; survivorship was zero in individuals homozygous for the reference allele (gray) and was 1 in individuals with at least one alternate allele (orange and purple). F: decay of linkage disequilibrium across the scaffold (blue points are means across 1000bp bins); LD remains high even at large distances between SNPs.

**Fig 3.**
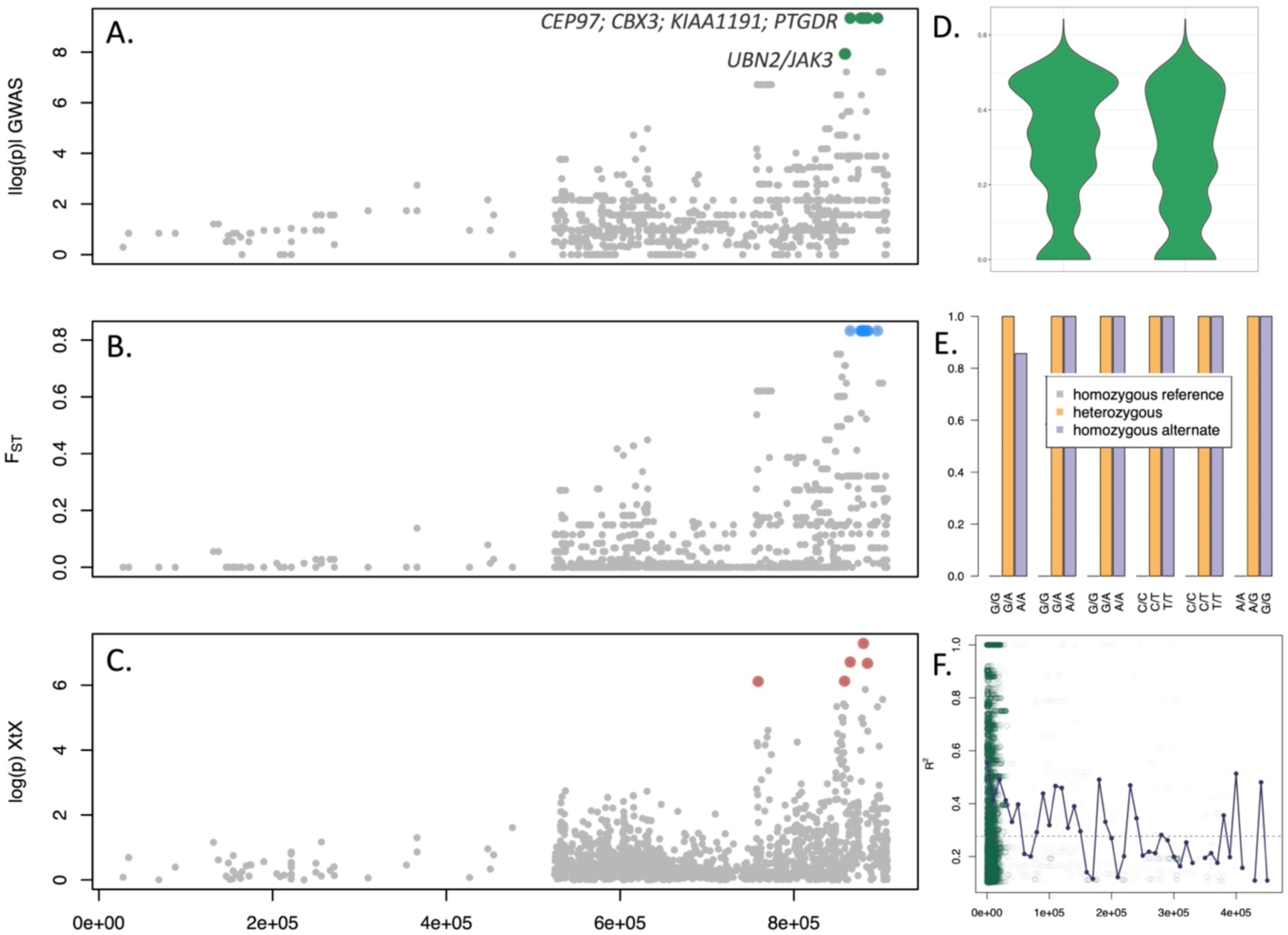
Candidate loci on Scaffold 519. A: -log(p) using genomic control adjusted p-values on a test in plink v1.9 for association between survivorship and genotype. B: F_ST_ between survivors and fatalities. C: -log(p) of the XtX selection statistic calculated in BayPass. D: Heterozygosity on this scaffold in fatalities (left) and survivors (right). E: Percent survivorship for each genotype at the six top candidate sites on scaffold 519; survivorship was zero in individuals homozygous for the reference allele (gray) and at ⅚ sites was 1 in individuals with at least one alternate allele (orange and purple). F: decay of linkage disequilibrium across the scaffold (blue points are means across 10,000bp bins); LD remains high even at large distances between SNPs.

### Candidate loci

Results from the three sets of analyses (GWAS, tests for selection in BayPass, and F_ST_) collectively resolved 56 candidate SNPs on 6 scaffolds that were common across multiple analyses and an additional 21 SNPs on three scaffolds that reached the highest significance in one model (Supplementary Table S4) and were characterized by alternate alleles in survivors only. Two SNPs were removed from the list of candidates because they were the only significant SNPs (even at the moderate threshold) on their scaffolds. We retained one SNP that was the only significant SNP on its scaffold at the stringent threshold because at the moderate threshold, 100 SNPs on the scaffold were significant. Therefore, 75 candidate SNPs were genotyped to assess whether survivors had alternate alleles (Supplementary Table S5–S6). Across 7 scaffolds, survivors had predominantly alternate alleles at 41 of these candidate SNPs (“top” and “secondary” candidates; Supplementary Table S7) while fatalities had primarily the reference allele, this pattern was particularly strong in 20 candidate SNPs across 5 scaffolds (“top candidates”, Table 2; Fig. 2-3; Supplementary Table S8; Supplementary Fig S3–S5). On some scaffolds (e.g., scaffold 443 and scaffold 519; Fig 2E and Fig 3E), survival was associated with either one or two alternate alleles at candidate SNPs, with no alternate alleles in fatalities, while on other scaffolds (e.g., scaffold 654; supplementary figure S5E), survivorship of individuals heterozygous at candidate SNPs was low, and survivorship probability was 1 only for individuals homozygous for the alternate allele. In yet other scaffolds (e.g., scaffold 375; supplementary figure S4E), the alternate allele was rarely found as a homozygote, but survivorship was high for heterozygotes, and fatalities had almost exclusively reference alleles. These differences highlight the different potential causative mechanisms by each SNP.

**Table 2.** List of top candidate sites and the top blast result for each.

| Scaffold | Positions | Inferred gene | Inferred gene function |
| --- | --- | --- | --- |
| scaffold22 | 221437, 221836, 225312 | cyclin A1 (*CCNA1*) (E=0) | Control of cell cycle |
| scaffold22 | 229648 | transmembrane protein 198 (*TMEM198*) (E=0) | positive regulation of canonical Wnt signaling pathway |
| scaffold22 | 235018 | CDK7 strongly-dependent group 2 enhancer (E=1e-142) | transcriptional regulatory region |
| scaffold22 | 236344 | cyclin A1 (*CCNA1*) (E=2e-67) | Control of cell cycle |
| scaffold375 | 202820 | 1. protocadherin beta-15 (*PCDHB15*) / beta-12 (*PCDHB12*) (E=0)<br>2. G protein-coupled receptor 65 (*GPR65*) (E=2e-134) | 1. integral plasma membrane proteins; play a role in the function of specific cell-cell neural connections<br>2. Plays a role in immune response by maintaining lysosome function and supporting phagocytosis-mediated intracellular bacteria clearance. May have a role in activation-induced cell death or differentiation of T-cells |
| scaffold375 | 218376 | Marmota monax G protein-coupled receptor 21 (*GPR21*) (E=0) | membrane proteins which activate signaling cascades as a response to extracellular stress |
| scaffold443 | 1030476, 1030693, 1058962 | ICOS gene for inducible T-cell costimulator protein (*ICOS*) (E=0) | Enhances all basic T-cell responses to a foreign antigen, namely proliferation, secretion of lymphokines, up-regulation of molecules that mediate cell-cell interaction, and effective help for antibody secretion by B-cells |
| scaffold519 | 859282 | 1. ubinuclein 2 (*UBN2*) (E=7e-60)<br>2. Janus kinase 3 (*JAK3*) (E=7e-50) | 1. involved in DNA replication-independent chromatin assembly; an important paralog of ubn1 which is downregulated in neutrophils confronted with *E. coli* and bacterial lipopolysaccharide (Zhang et al. 2004)<br>2. Mediates essential signaling events in both innate and adaptive immunity and plays a crucial role in hematopoiesis during T-cells development |
| scaffold519 | 864821 | centrosomal protein 97 (*CEP97*) (E=1e-154) | negative regulation of cilium assembly and regulation of mitotic spindle assembly |
| scaffold519 | 877694, 878376 | chromobox 3 (*CBX3*) (E=2e-74) | involved in transcriptional silencing in heterochromatin-like complexes |
| scaffold519 | 880072 | KIAA1191 ortholog (*KIAA1191*) (E=0) | enable oxidoreductase activity |
| scaffold519 | 884668 | prostaglandin D2 receptor (*PTGDR*) (E=3e-125) | transmembrane proteins that respond to extracellular cues and activate intracellular signal transduction pathways. PGD2 involved in mediating inflammation. |
| scaffold654 | 334941, 354763, 364656 | succinyl-CoA:glutarate-CoA transferase (*SUGCT*) (E=0) | Catalyzes the succinyl-CoA-dependent conversion of glutarate to glutaryl-CoA |

### Candidate Gene Identification

For our 20 top candidate SNPs and 21 secondary candidates, our BLAST search resolved candidate genes in at least one, and sometimes multiple, hits per locus (Table 2, Supplementary Table S9). Some of these genes’ primary role is in the immune system, such as inducible T-cell costimulator protein (*ICOS*; scaffold 443) and selection and upkeep of intraepithelial T-cells protein 8 (*SKINT8*; scaffold 611). In addition, several candidate genes act directly to regulate immune processes, including prostaglandin D2 receptor (*PTGDR*; scaffold 519), which is involved in mediating inflammation, chromobox 3 (*CBX3*, scaffold 519), which can upregulate interleukin 1A (M. Yang et al. 2022), and eukaryotic translation initiation factor 2 alpha kinase 2 (*EIF2AK2*, scaffold 611), a gene in the TLR pathway that plays a regulatory role in the expression of genes encoding pro-inflammatory cytokines and interferons (Hales 2021). Finally, there were several candidate genes for which we were unable to find prior evidence of involvement in combating infection but that may have an as-yet undetermined role in immunity (Table 2, Supplementary Table S9).

## Discussion

In a world where globalization is drastically increasing the number of introduced pathogens, there is an urgent need to understand the potential for rapid adaptation in immunologically naive species. Pathogens can be especially destructive in keystone species such as prairie dogs, on which numerous other species--including the endangered black-footed ferret--rely for food, shelter, and defense against predators. We capitalized on one of the only documented cases of survival during an epizootic of sylvatic plague in prairie dogs to assess the genomic basis of evolved response to an introduced virulent pathogen. Genomic signatures of adaptation were apparent in surviving prairie dogs, with evidence of selection especially strong across five scaffolds. These scaffolds contain regions that blast to both immune and non-immune genes. We infer that rapid adaptation to plague is likely multigenic, with a primary role of T-cell related genes.

Our knowledge of the interaction between *Y. pestis* and hosts at the early stages of infection is limited (R. Yang et al. 2023). Although the mechanistic action of resistance is unknown, this work identified several new candidate genes whose role in adaptation to *Y. pestis* has not been demonstrated (e.g., cyclin A1 (*CCNA1*; scaffold 22) and succinyl-CoA:glutarate-CoA transferase (*SUGCT*; scaffold 654)), and it also provided support for genes with previously identified or hypothesized roles in infections with bacterial pathogens (e.g., chromobox 3 (*CBX3*; scaffold 519) upregulates interleukin 1A (M. Yang et al. 2022); G protein-coupled receptor 21 (*GPR21*; scaffold 519) and 65 (*GPR65*; scaffold 519) are membrane proteins that respond to extracellular stress, mediate inflammation and support lysosome function and bacterial phagocytosis (Tan et al. 2017; Mercier et al. 2022); and prostaglandin D2 receptor (*PTGDR*; scaffold 519) mediates inflammation (Murata et al. 2013; Sheppe and Edelmann 2021), including in infection with *Y. pestis* (Brady et al. 2024)). Of particular interest is the inducible T-cell stimulator gene (*ICOS*; scaffold 443), a gene that enhances T-cell responses to foreign antigens (Dong et al. 2001) and which was also found to be under selection in humans during the Black Death (Klunk et al. 2022). Interestingly, three candidate genes identified here (*TMEM198*, *PCDHB12/15*, and *KIAA1191*) are different from but in the same gene classes (transmembrane proteins, protocadherins, and Kasuza protein binding genes) as candidate genes for plague resistance in great gerbils (*TMEM200A*, *PCDHB18*, and *KIAA0408*) (Nilsson et al. 2022). This finding provides support for the hypothesis that parallel evolution may occur at the level of gene classes in addition to individual genes (Cassin-Sackett, Callicrate, and Fleischer 2019).

The limited number of survivors in an epizootic in a natural population leads to statistical challenges in identifying the genomic basis of complex traits, including the risk of both Type I and Type II errors. We aimed to minimize the number of false positives by focusing only on sites that were significant in multiple tests. Similar approaches have been used with success previously (Cornetti and Tschirren 2020), but the cost of this approach is the inflated rate of false negatives (Rice, Schork, and Rao 2008) (an effect that is amplified in small sample sizes), such that we are likely to detect only loci of large effect (Ioannidis, Tarone, and McLaughlin 2011). Nonetheless, this tradeoff may be appropriate in the context of designing genetically informed management actions, which are likely to prioritize such large-effect loci. study offers three important insights. First, some secondary candidate sites (e.g., those on scaffold 375, 611, and 1325; Supplementary Fig. S4) show a single alternate allele in each survivor and none in fatalities, suggesting these could be recent mutations that contributed to survivorship (Peter, Huerta-Sanchez, and Nielsen 2012). Other candidate sites (e.g., those on scaffolds 443, 519, and 654; Figs 2e, 3e and Supplementary Fig. S5) are characterized by near-fixation of the alternate allele in the population of survivors and 1 - 2 alternate alleles in the population of fatalities, consistent with predictions of directional selection on standing variation (McManus et al. 2017; Dionne et al. 2009). Even with the low number of survivors in this epizootic, the approach of using multiple tests resolved many of the same candidate sites across tests (Tables S2–S3), including sites with these contrasting evolutionary histories (i.e., recent mutations versus standing variation).

Second, recently evolved immunity may be conferred by a combination of immune and non-immune genes (Elfekih et al. 2022; Cassin-Sackett, Callicrate, and Fleischer 2019), including gene duplication and specialization of immune genes (Nilsson et al. 2020), with non-immune genes likely to exhibit pleiotropic effects in combating infection with a novel pathogen. In genes for which pleiotropy is antagonistic, as in the genes causing sickle-cell anemia in some malaria-resistant human individuals (Grosse et al. 2011; Elguero et al. 2015), we may predict slower rates of adaptation at these loci due to costs of resistance. This may be why prairie dog populations have been extirpated even in places where plague survivors had previously been observed (e.g., Gunnison’s prairie dogs in northern New Mexico (Cully et al. 1997)).

Third, subpopulations within a metapopulation share genotypes for resistance-associated SNPs, potentially indicating the recent spread of adaptive alleles (Gignoux-Wolfsohn et al. 2021). Moreover, differentiation among subpopulations was dramatically lower at candidate than neutral sites, suggesting heightened rates of gene flow of adaptive variation (Matz et al. 2018) even in urbanized landscapes. Adaptive variants may percolate through the landscape at a faster rate than neutral variants, following pathways of connectivity through a complex landscape matrix (Sackett et al. 2012).

Nonetheless, the extent to which these variants persist in natural populations is currently unknown. Although there has been extremely strong selection for resistance for 25-50 prairie dog generations (Adjemian et al. 2007), selection from plague is intermittent, and this temporal variation in selection means that individuals in one generation may not be exposed to selection. This fluctuating selection may act to maintain adaptive genetic variants in a population but without leading to fixation (O. L. Johnson et al. 2023). Conversely, adaptive variants may be lost by drift or masked by gene dilution in the absence of selection; indeed, some populations that were spared in a 1994 epizootic (Collinge et al. 2005) were extirpated in the 2006 epizootic (Sackett, Collinge, and Martin 2013b). Long-term effects on allele frequencies thus hinge on selective and neutral processes occurring in non-plague years in addition to the years in which strong selection for resistance is exerted (Lynch et al. 2024). Moreover, selection may be counteracted by fitness costs of resistance—particularly if survival is conferred by pleiotropic effects of non-immune genes (Taylor, Cerami, and Fairhurst 2013; Byars and Voskarides 2020). Quantifying long-term patterns in allele frequencies at these candidate sites—an endeavor that is increasingly possible in species with consistent museum collection records (Byerly et al. 2024)—could help shed light on the degree to which the long-term persistence of adaptation is possible via natural or facilitated mechanisms.

The evolutionary outcome of strong selection followed by the emergence of resistance to sylvatic plague in prairie dogs is still unclear, and data from experimental infections (Rocke et al. 2012; Busch et al. 2013; Rocke et al. 2015) are needed to understand mechanistic relationships between genotype and phenotype. Yet our results demonstrate that there is genetic variation within populations for resistance to a recently introduced highly virulent pathogen, and that strong associations exist between genotype and phenotype at several loci. The relatively small number of loci associated with recently evolved resistance include a mix of classic immune and non-immune genes—one of which has been associated with survival from plague in humans (Klunk et al. 2022). Others are related to genes hypothesized to combat plague in other rodents (Nilsson et al. 2022), suggesting that rapid adaptation can occur when there is sufficient standing variation in genes involved in pathways commonly invoked to fight pathogens. The existence of multiple genes conferring survival across taxa suggests that native species have the potential for adaptation to introduced pathogens—an increasingly important concern in an unprecedented era of emerging infectious diseases.

Understanding the genetic basis of adaptation is a first step in effective long-term management of disease risk in threatened and endangered species, as it can inform actions such as facilitated adaptation (Torres et al. 2023) via relocation of resistant populations to priority conservation areas. However, because adaptation occurs over multiple generations and is often reliant on existing standing variation, conservation stopgaps are also needed. In the U.S., land managers prevented the extinction of the then-critically endangered black-footed ferret in part by developing a plague vaccine for prairie dogs (Rocke et al. 2008), which increases survivorship probability during epizootics (Rocke et al. 2017). This vaccine and other interventions such as control of the flea vectors (tripp, eads papers) has enabled specific prairie dog populations to persist in the short term while the incidence of natural immunity increases across the species range (Rocke et al. 2012) due to selection on adaptive variants. Given the enormous financial investment in protecting endangered species, it would be useful for managers to know the general time frame required for the evolution of immunity to emerging pathogens, and to use this information in conservation planning. Our results suggest that, in species with high levels of standing variation (Vonhof, Russell, and Miller-Butterworth 2015) (e.g., a variable site every 400 bp such as observed in the 14 sampled individuals here) and strong selection, we may expect immunity to evolve within ∼25 generations. In pathogens with lower virulence (Kollias et al. 2004), selection is weaker and survivorship is higher, likely accompanied by the slower evolution of immunity (except in species with very high effective population sizes (Bonneaud et al. 2011; Henschen et al. 2023)) but necessitating fewer conservation interventions. Conversely, in species with smaller effective population sizes (Cassin-Sackett et al. 2021), conservation interventions will likely be needed long-term to prevent extinction.

Ultimately, understanding the genetic basis of immunity-associated alleles will pave the way for other intriguing questions, including: i) Can we predict the rate of host evolution of immunity to a novel pathogen given measurable parameters such as pathogen virulence and the degree of standing variation in the host? ii) Is there a quantifiable minimum effective population size required not only to prevent inbreeding, but also to enable adaptation to the ongoing changes in the Anthropocene? iii) What is the optimum amount of gene flow through an urban or natural landscape that optimizes the spread of adaptive alleles while preventing their dilution and loss, particularly when selection is episodic? The system of recently evolved immunity to plague in prairie dogs is poised to become a powerful model for conservation planning of threatened and endangered species.

## Acknowledgments

The individual sampled for the genome assembly was sampled under Colorado Parks and Wildlife scientific collection license #17TR2012a. Funding for sample collection was provided by the University of Colorado (CU), CU’s Department of Ecology and Evolutionary Biology, the CU Museum of Natural History, and the Boulder County Nature Association. Other support was provided by the University of South Florida and NSF IOS-2220815. Computing was conducted on the Smithsonian Institution High Performance Cluster (SI/HPC), Smithsonian Institution (https://doi.org/10.25572/SIHPC) and on the Louisiana Optical Network Infrastructure (LONI, https://loni.org/). We are grateful to Erin Arnold and other field assistants who helped sample prairie dogs. Boris Schmid provided insightful comments that improved this manuscript. We would like to thank Boulder Open Space and Mountain Parks, Boulder County Open Space, and Boulder Parks and Recreation for permission to live-trap prairie dogs on public lands. Sampled colonies were located on unceded Arapaho, Cheyenne, Núu-agha-tv-p, and Oceti Sakówin lands.

