## Supplementary Figure for "Rapid adaptation to a globally introduced virulent pathogen in a keystone species"

### Supplementary Methods

*Genome Annotation*

The genome was annotated as in (Tsuchiya et al. 2020). Briefly, we used RepeatMasker (open-4.0.6, Smit et al. 2013–2015) with the Rodentia database to identify repetitive elements in the genome and soft-mask the assembly. Next, we used BLAT (Kent 2002) to align our masked genome assembly to the proteins of both the closely related *Ictidomys tridecemlineatus* and the model organism *Mus musculus*. We used the dnax option to translate the DNA sequences to protein in six frames. Next, we generated a hints file for AUGUSTUS from three lines of evidence: 1) BLAT results, 2) BUSCO training parameters, and 3) masking information from RepeatMasker. To speed up the analysis, we partitioned our assembly into scaffolds (Haas et al. 2008) and ran AUGUSTUS in each scaffold individually. We extracted both the protein and nucleotide sequences of the gene models, as well as the individual coding sequences, in AUGUSTUS. Finally, we ran Blast (v2.6.0þ, Altschul et al. 1990) on the gene models identified by AUGUSTUS and used the final .xml file as an input to Blast2GO (v5.2.5, Götz et al. 2008) to functionally annotate the genome.

*Tests for outliers in OutFLANK*

We thinned the SNP dataset in the bigsnpr package for R using a r2 threshold of 0.3 and then executed OutFLANK v0.2 using a LeftTrimFraction and RightTrimFraction of 0.05, Hmin 0.1, and q threshold of 0.05, and limiting the dataset to sites with a minor allele count of at least 3. We also explored other values of RightTrimFraction and Hmin, but these did not change the number of inferred outliers.

*Heterozygosity*

We used bash scripts to count the number of heterozygotes and homozygotes in 100k bp bins and calculate a heterozygote ratio for each individual. We used an unpaired t-test in R v4.3.0 (R Core Team 2023) to test for differences in heterozygote ratio between survivors and fatalities and examined plots of scaffolds containing candidate SNPs to determine whether there were regions of low homozygosity in survivors that could be indicative of selective sweeps.

### Supplementary Results

*Tests for outliers in OutFLANK*

With a q threshold of 0.05, OutFLANK detected 1637 outliers between survivors and fatalities (at q=0.045, 1602 outliers were detected and at q=0.04 and below, no outliers were detected). When we altered the parameters of dataset thinning or the OutFLANK function (e.g., using a higher r2 threshold, lower LeftTrimFraction, etc.), we did not gain any additional resolution; the same 1602 outliers as above were recovered under different scenarios. 88 SNPs had both raw and corrected FST estimates of 1, but were statistically indistinguishable from sites with much lower FST, suggesting that lack of resolution -- i.e., the inability to recover fewer loci with a more stringent q threshold-- stemmed from a lack of statistical power rather than a lack of SNPs that were highly differentiated between survivors and fatalities.

*Heterozygosity*

At the level of scaffolds, 3 of the 9 scaffolds with candidate SNPs showed significant differences in heterozygote ratio between survivors and fatalities (unadjusted $p < 0.001$), with heterozygote ratio being higher in survivors on scaffolds 22 and 375, and lower in survivors on scaffold 377. Several of the scaffolds with candidate SNPs exhibited reduced heterozygosity in survivors in certain regions only.

### Supplementary Figures

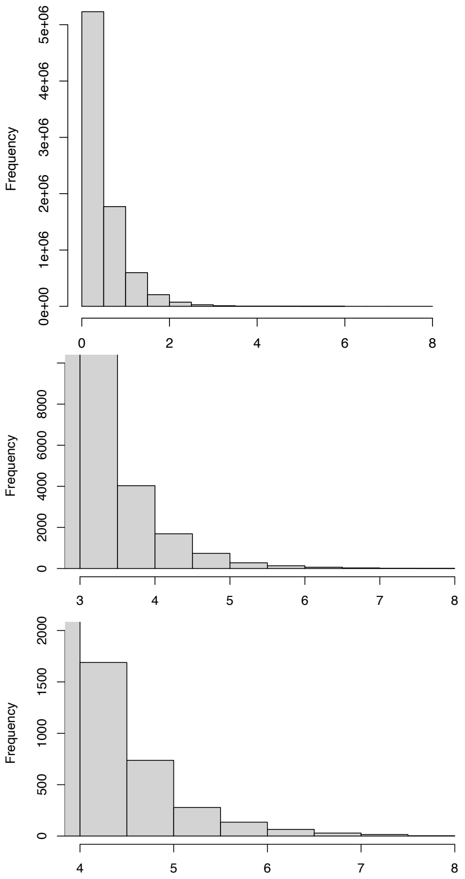


**Supplementary Figure 1**. Distribution of -log(10) p values from BayPass. Top panel: raw histogram, middle panel: zoomed-in region of -log(10) p > 3, and bottom panel: zoomed-in region of -log(10) p > 4.

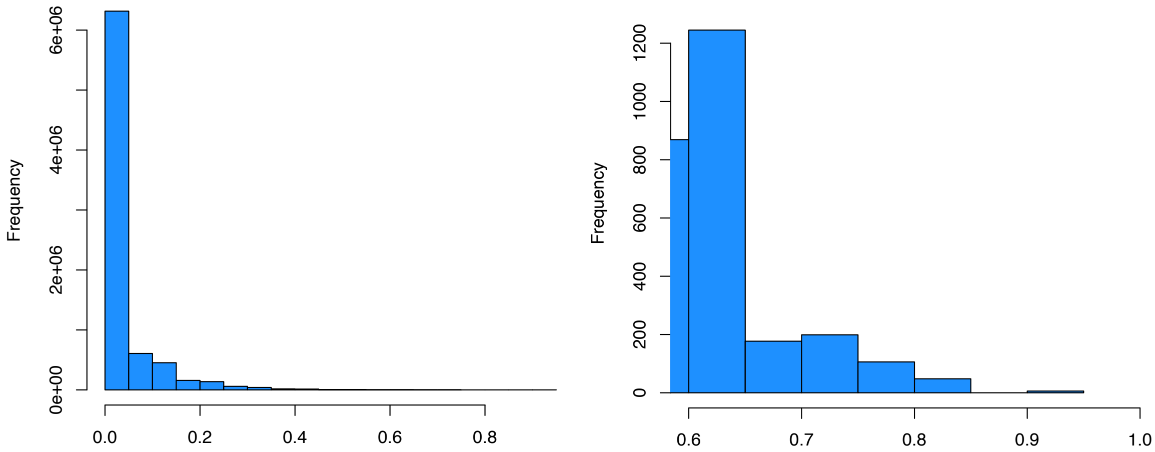


**Supplementary Figure 2**. Distribution of $F_{ST}$ values between survivors and fatalities. Left panel: all $F_{ST}$ values calculated with the Weir & Cockerham method in VCFtools, with negative values converted to zeroes. Right panel: zoomed-in region of $F_{ST}$ > 0.6.

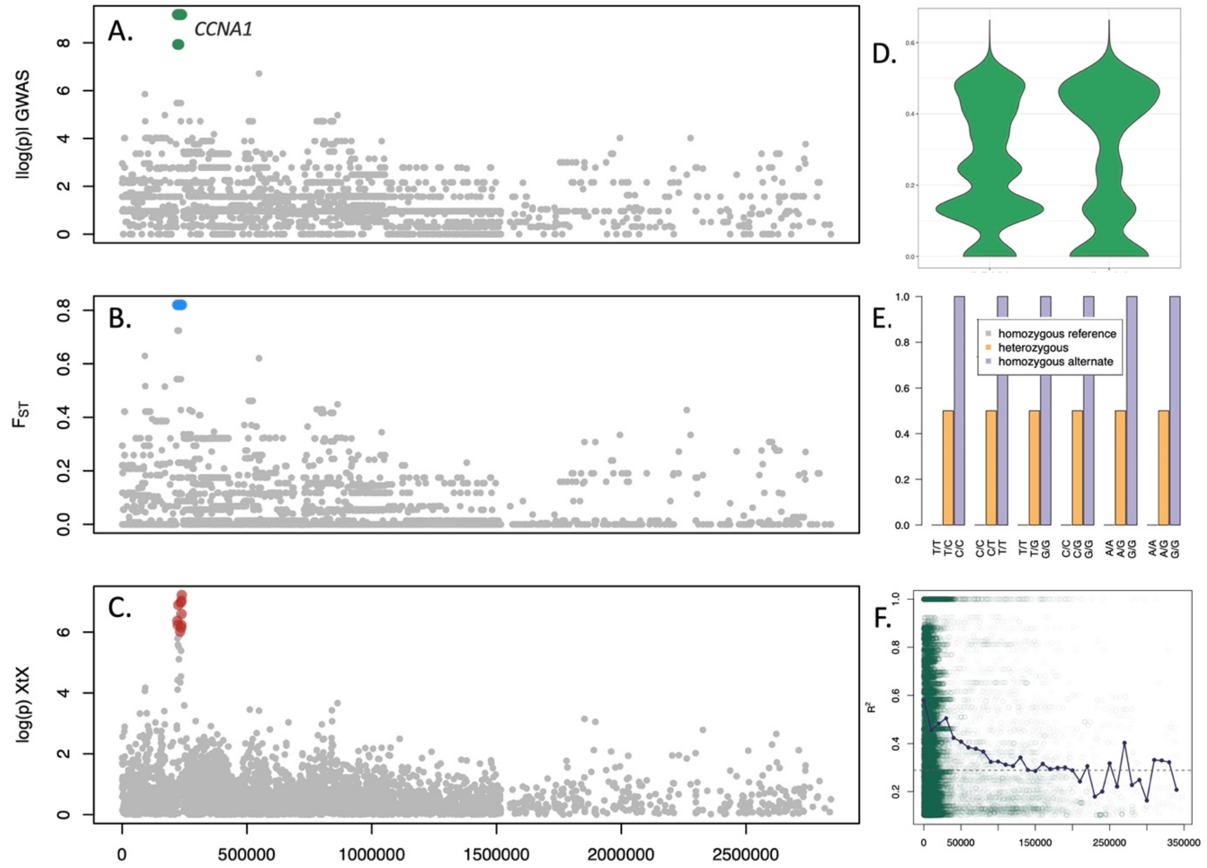


**Supplementary Figure 3**. Candidate loci on Scaffold 22. A: -log(p) using genomic control adjusted p-values on the test in plink v1.9 for association between survivorship and genotype. B: $F_{ST}$ between survivors and fatalities. C: -log(p) of the XtX selection statistic calculated in BayPass. D: Heterozygosity on this scaffold in fatalities (left) and survivors (right). E: Percent survivorship for each genotype at the six top candidate sites on scaffold 22; percent survivorship was zero in individuals homozygous for the reference allele (gray), intermediate (0.5) in heterozygotes (orange), and 1 in individuals homozygous for the alternate allele (purple). F: decay of linkage disequilibrium across the scaffold (blue points are means across 1000bp bins); LD remains high even at large distances between SNPs.

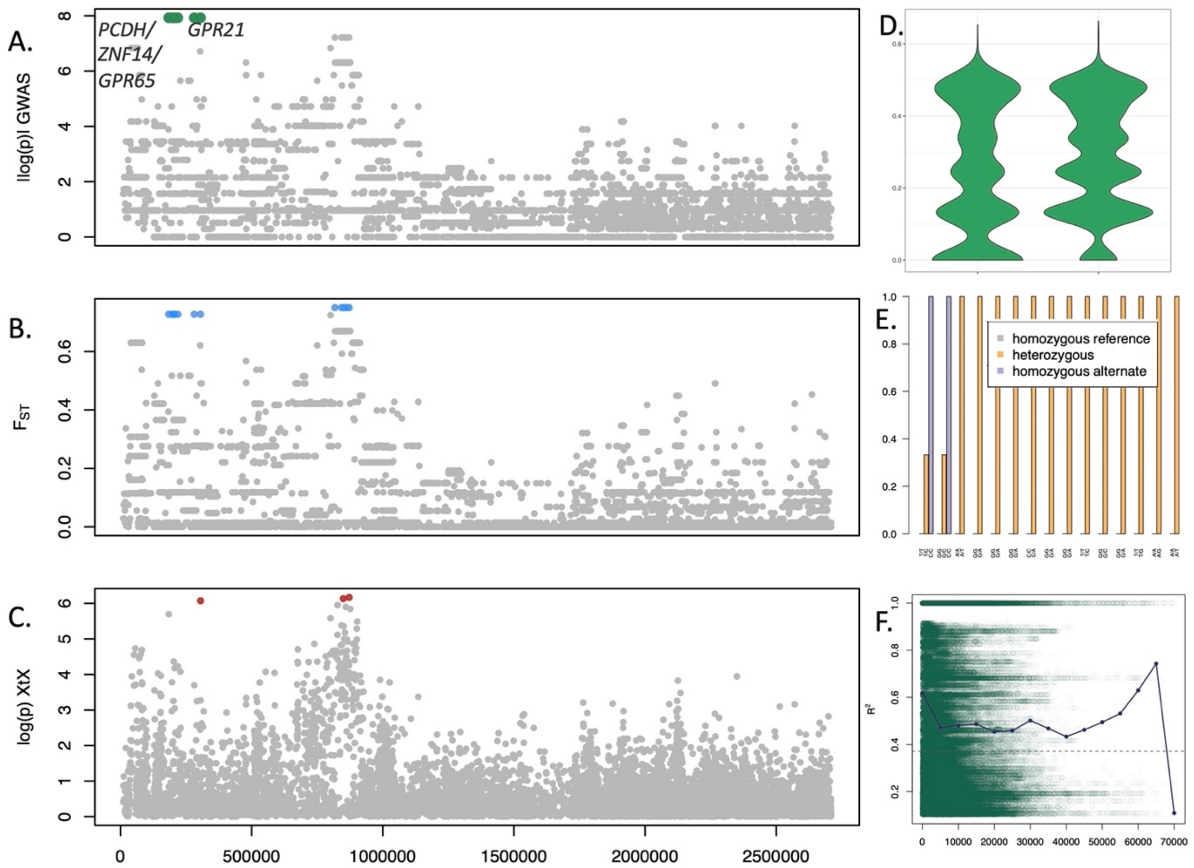


**Supplementary Figure 4**. Candidate loci on Scaffold 375. A: -log(p) using genomic control adjusted p-values on the test in plink v1.9 for association between survivorship and genotype. B: $F_{ST}$ between survivors and fatalities. C: -log(p) of the XtX selection statistic calculated in BayPass. D: Heterozygosity on this scaffold in fatalities (left) and survivors (right). E: Percent survivorship for each genotype at the three top candidate sites on scaffold 375; at the top two sites survivorship was zero in individuals homozygous for the reference allele (gray), intermediate (0.33) in heterozygotes (orange), and 1 in individuals homozygous for the alternate allele (purple). At the secondary candidate sites, survivorship was zero in individuals homozygous for the reference allele (gray) and 1 in heterozygotes (orange); homozygotes for the alternate allele (purple) were not observed. F: decay of linkage disequilibrium across the scaffold (blue points are means across 1000bp bins); LD remains high even at large distances between SNPs. Note: y-axis scale is different in parts B and C than figures in the main text, and SNPs at the moderate significance threshold are highlighted.

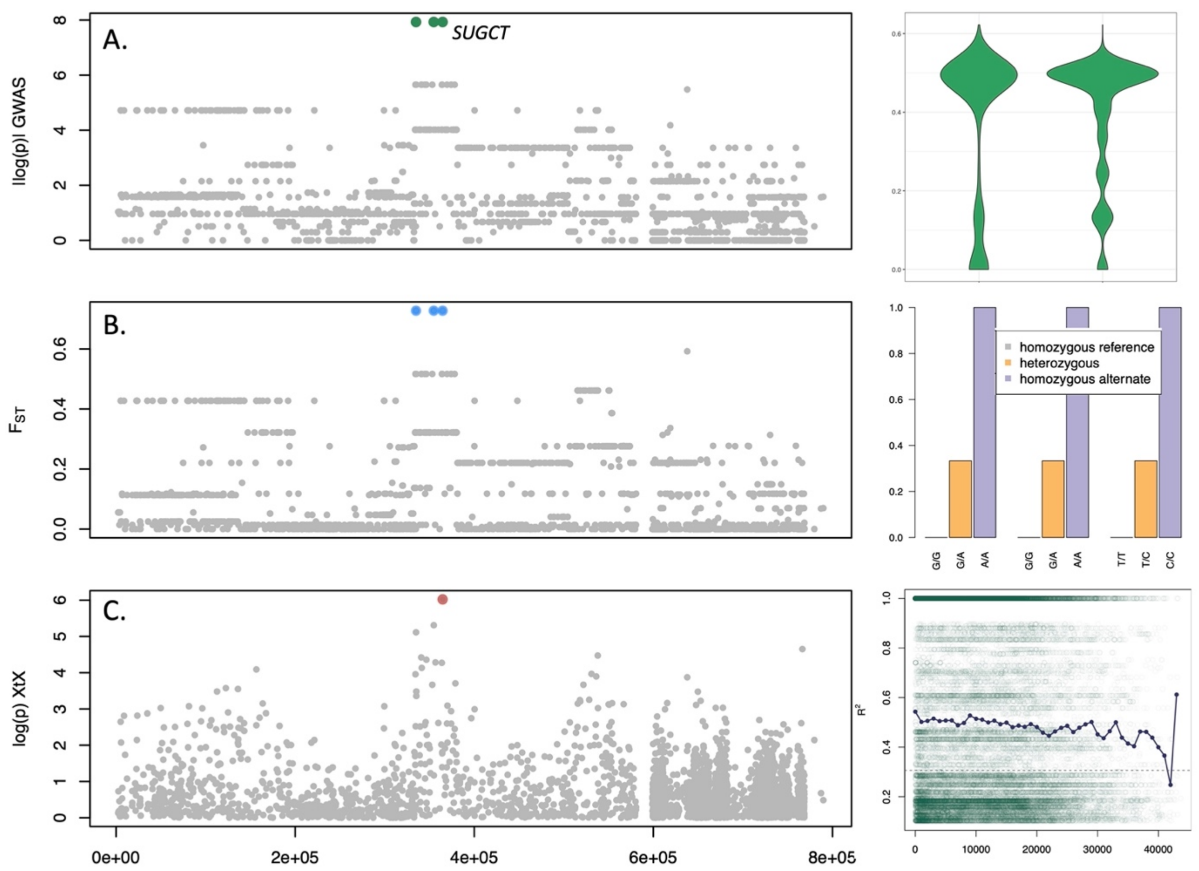


**Supplementary Figure 5**. Candidate loci on Scaffold 654. A: -log(p) using genomic control adjusted p-values on the test in plink v1.9 for association between survivorship and genotype. B: $F_{ST}$ between survivors and fatalities. C: -log(p) of the XtX selection statistic calculated in BayPass. D: Heterozygosity on this scaffold in fatalities (left) and survivors (right). E: Percent survivorship for each genotype at the three top candidate sites on scaffold 654; survivorship was zero in individuals homozygous for the reference allele (gray), intermediate (0.33) in heterozygotes (orange), and 1 in individuals homozygous for the alternate allele (purple). F: decay of linkage disequilibrium across the scaffold (blue points are means across 1000bp bins); LD remains high even at large distances between SNPs. Note: y-axis scale is different in parts B and C than figures in the main text, and SNPs at the moderate significance threshold are highlighted.

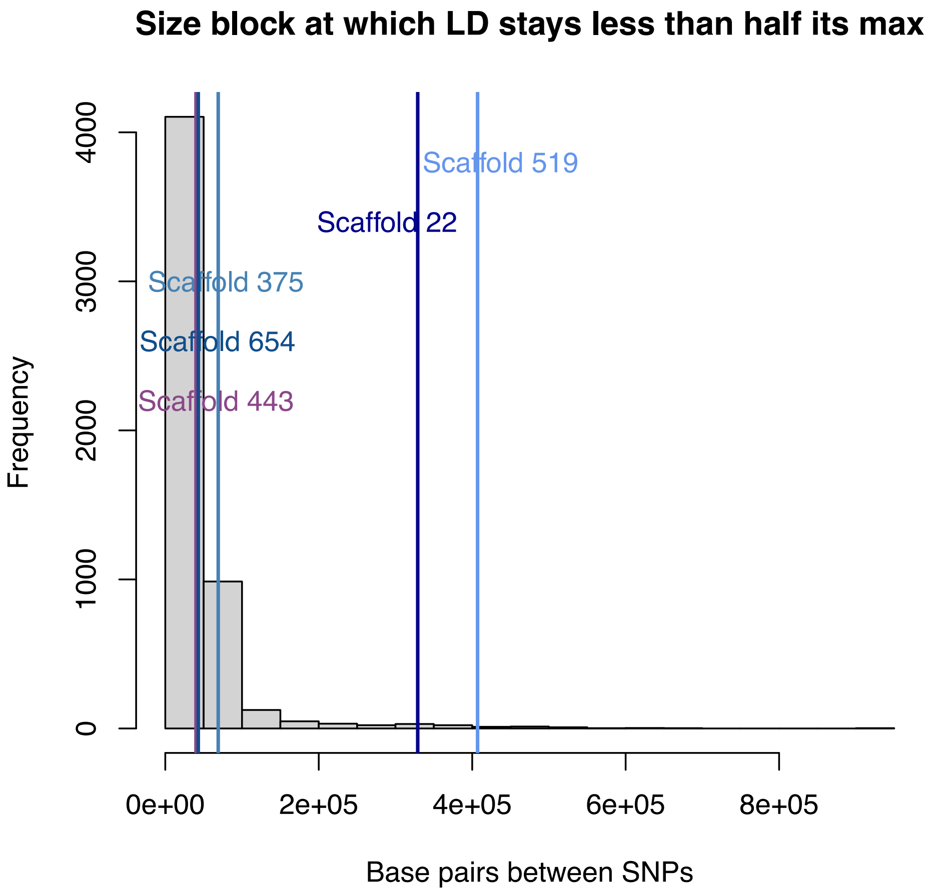


**Supplementary Figure 6**. Distribution across all scaffolds of the size of LD blocks at the scaffold's LD50. Many scaffolds had a max $R^2$ of 1, so this often marks the size of LD blocks with $R^2$=0.5. Scaffolds with candidate SNPs had larger than average blocks of LD with $R^2$ at half its max, with scaffold22 and scaffold519 retaining particularly large blocks of high LD.

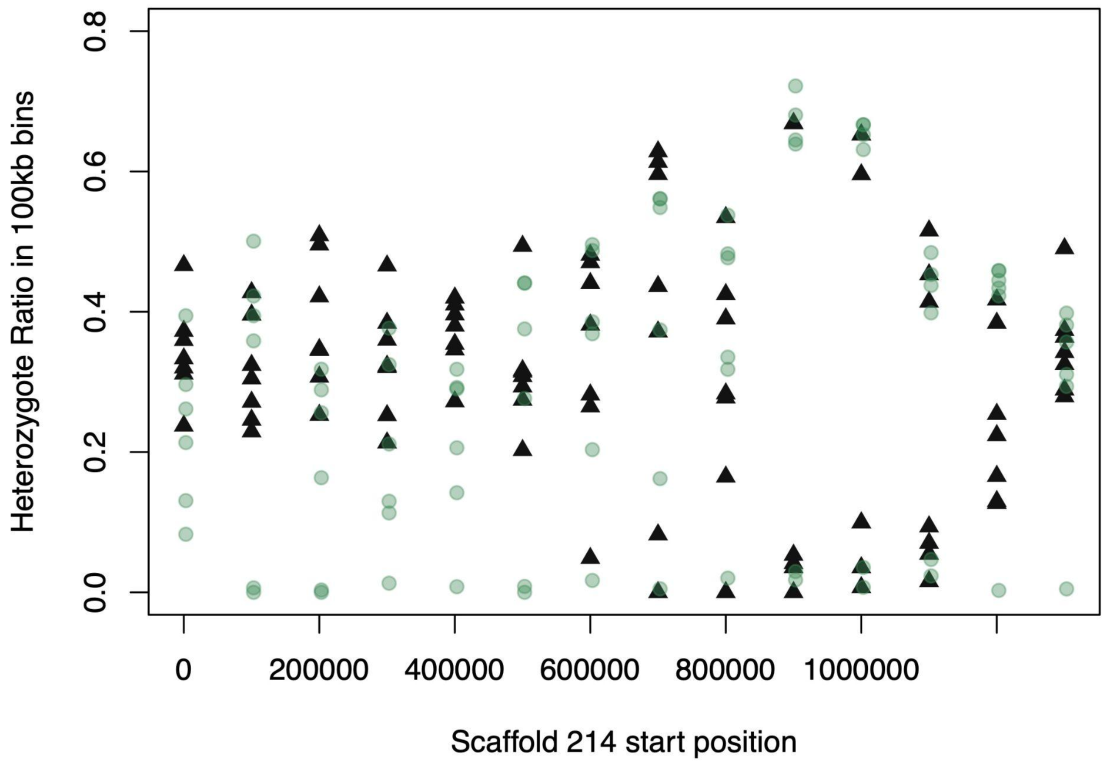


Fig x. Proportion of total sites that were heterozygous in survivors (green circles) or fatalities (black triangles) along scaffold 214. In the first half of the scaffold, heterozygosity is lower in survivors.
